# Spatial inhibition of return is impaired in mild cognitive impairment and mild Alzheimer’s disease

**DOI:** 10.1101/2020.05.11.089383

**Authors:** Xiong Jiang, James H. Howard, G. Wiliam Rebeck, R. Scott Turner

## Abstract

Spatial inhibition of return (IOR) refers to the phenomenon by which individuals are slower to respond to stimuli appearing at a previously cued location compared to un-cued locations. Here we provide evidence supporting that spatial IOR is mildly impaired in individuals with mild cognitive impairment (MCI) or mild Alzheimer’s disease (AD), and the impairment is readily detectable using a novel double cue paradigm. Furthermore, reduced spatial IOR in high-risk healthy older individuals is associated with reduced memory and other neurocognitive task performance, suggesting that the novel double cue spatial IOR paradigm may be useful in detecting MCI and early AD.

**SIGNIFICANCE STATEMENT:**

- Novel double cue spatial inhibition of return (IOR) paradigm revealed a robust effect IOR deficits in individuals with mild cognitive impairment (MCI) or mild Alzheimer’s disease (AD)
- Spatial IOR effect correlates with memory performance in healthy older adults at a elevated risk of Alzheimer’s disease (with a family history or APOE e4 allele)
- The data suggests that double cue spatial IOR may be sensitive to detect early AD pathological changes, which may be linked to disease progress at the posterior brain regions (rather than the medial temporal lobe)

## INTRODUCTION

Spatial inhibition of return (IOR) refers to the phenomenon by which individuals are slower to respond to stimuli appearing at a previously cued location compared to un-cued locations ^1^. In a classic spatial cue-target paradigm, subjects are usually faster to respond to the target appearing at the cued than un-cued location when the stimuli onset asynchrony (SOA) between the target and cue is short (∼200ms or less), but are slower when the SOA is long (∼300-500ms or more), with the latter generally referred as spatial IOR. First reported in 1984 ^2^, spatial IOR has been studied using different modalities of stimuli ^3^, different responses (i.e., manual vs. saccadic response) ^4,5^, and in older adults ^6^.

The temporoparietal junction (TPJ) and the inferior parietal cortex have been shown to play critical roles in spatial IOR ^1^. Given that these regions are implicated in AD progression ^7^, this suggests that spatial IOR may be useful in MCI and AD diagnosis. However, it remains controversial whether spatial IOR is impaired in AD. Early studies suggest that spatial IOR is relatively preserved in AD ^8–10^. By contrast, more recent studies suggest that spatial IOR may be impaired in individuals with MCI or AD ^11–13^, and IOR deficits in individuals with MCI may be predictive of conversion to dementia ^13^. Most previous studies employed the classic single cue-target paradigm ^2^.

Previous studies have suggested that increasing the number of cues may lead to an increase in IOR effect ^14^. Here we investigated probable spatial IOR deficits in Alzheimer’s disease using a modified double cue spatial IOR task.

## MATERIALS AND METHODS

### Participants

Eight individuals with MCI, eight individuals with mild AD, and 41 healthy older adults participated in the study (Table 1). Prior to enrollment, a signed informed consent form approved by the Georgetown University Medical Center’s Institutional Review Board was obtained from all participants and their legally authorized representatives (if they had a diagnosis of MCI or mild AD). Data from a few additional participants were excluded from the analysis due to various reasons (Supplementary Materials), but the data from these excluded subjects is shown in Supplementary Fig. 1.

**Table 1.** The demographics and neuropsychological test scores of control, MCI, and mild AD participants. ADAS-Cog, Alzheimer’s Disease Assessment Scale-Cognitive subscale; CA, Caucasian-Americans; LADL, Lawton Instrumental Activities of Daily Living Scale; LM, Logical Memory Test; LVF, Letter Verbal Fluency; MCI, mild cognitive impairment; MMSE, mini-mental state exam; MoCA, Montreal Cognitive Assessment; NPI, Neuropsychiatric Inventory.

|  | **Controls** | **MCI/AD** | ***p*<sup>2</sup>** |
| --- | --- | --- | --- |
| N (F) | 41 (26F<sup>1</sup>) | 16 (5F<sup>1</sup>) | 0.040<sup>3</sup> |
| Age | 67.4 ± 5.3 | 69.3 ± 4.8 | n.s. |
| Education (yrs) | 18.4 ± 3.4 | 19.5 ± 4.2 | n.s. |
| %CA | 78.1% | 86.% | n.s.<sup>3</sup> |
| APOE ε4 carriers (%) | 29.3% | 60.0% | 0.028<sup>3</sup> |
| AD family history (%) | 36.6% | 46.7% | n.s.<sup>3</sup> |
| MMSE | 29.3 ± 1.0 | 27.6 ± 2.0 | 0.0001 |
| MoCA<sup>4</sup> | 25.2 ± 1.8 | 21.4 ± 4.4 | 0.0089 |
| LM Immediate | 12.0 ± 3.5 | 8.3 ± 3.9 | 0.0011 |
| LM Delayed | 9.5 ± 4.2 | 5.7 ± 4.3 | 0.0045 |
| ADAS-cog | 6.3 ± 3.4 | 14.5 ± 6.1 | 5.0E-08 |
| NPI | 2.3 ± 4.9 | 3.9 ± 4.6<sup>5</sup> | n.s. |
| LADL | 76.3 ± 3.0 | 71.9 ± 8.2<sup>5</sup> | 0.0054 |
| LVF | 48.2 ± 13.1 | 41.2 ± 13.0 | n.s. |

<sup>1</sup> female, adding sex as a covariate produced similar (nearly identical) results; <sup>2</sup> uncorrected *p* values for the difference between controls and MCI/AD with two-tailed two-sample *t*-tests (unless otherwise specified); <sup>3</sup> Fisher's Exact Test; <sup>4</sup> MoCA were only administered to a subset of participants, including 17 controls and 10 MCI/AD patients (out of 41 controls and 15 MCI/AD patients included here; <sup>5</sup> NPI and LADL is no available in one MCI participant.

### Neuropsychological and other assessments

The following data were collected from all participants: blood pressure; biographical and health questionnaire; family history of AD; Mini-Mental State Examination (MMSE); Montreal Cognitive Assessment (MoCA); Alzheimer’s Disease Assessment Scale – Cognitive subscale (ADAS-Cog); Lawton Instrumental Activities of Daily Living Scale (LADL); Neuropsychiatric Inventory (NPI); Letter Verbal Fluency (LVF); Logical Memory subtest of the Wechsler Memory Scale (WMS) – fourth edition (WMS-IV). Saliva samples were collected from all except one AD patient for APOE genotyping.

### Spatial IOR Experimental Design

The experimental paradigm was adapted from a previous learning study ^15^. Within each trial, three stimuli were presented sequentially, two cue stimuli (solid red squares) and one target stimulus (solid green square). The subjects were instructed to observe the two red cues and respond to the green target to indicate by pressing one of two buttons in the right hand (with the index and the middle finger) to indicate whether the green target was presented at the left or right location (Fig. 1A). The two cue stimuli could appear for 200ms each in any of the three possible locations (left, center, right, shown as the three empty squares in Fig. 1A), whereas the target stimuli could only appear in two possible locations (left or right, but not center) for 850ms, then was followed by a 750ms blank screen before next trial started. Subjects had to respond within the 1.6sec time-window. Two runs of data were collected from all except 3 subjects, who only finished one run of experiment. There were 130 trials per run, and each trial lasted 2.5s.

**Figure 1.**
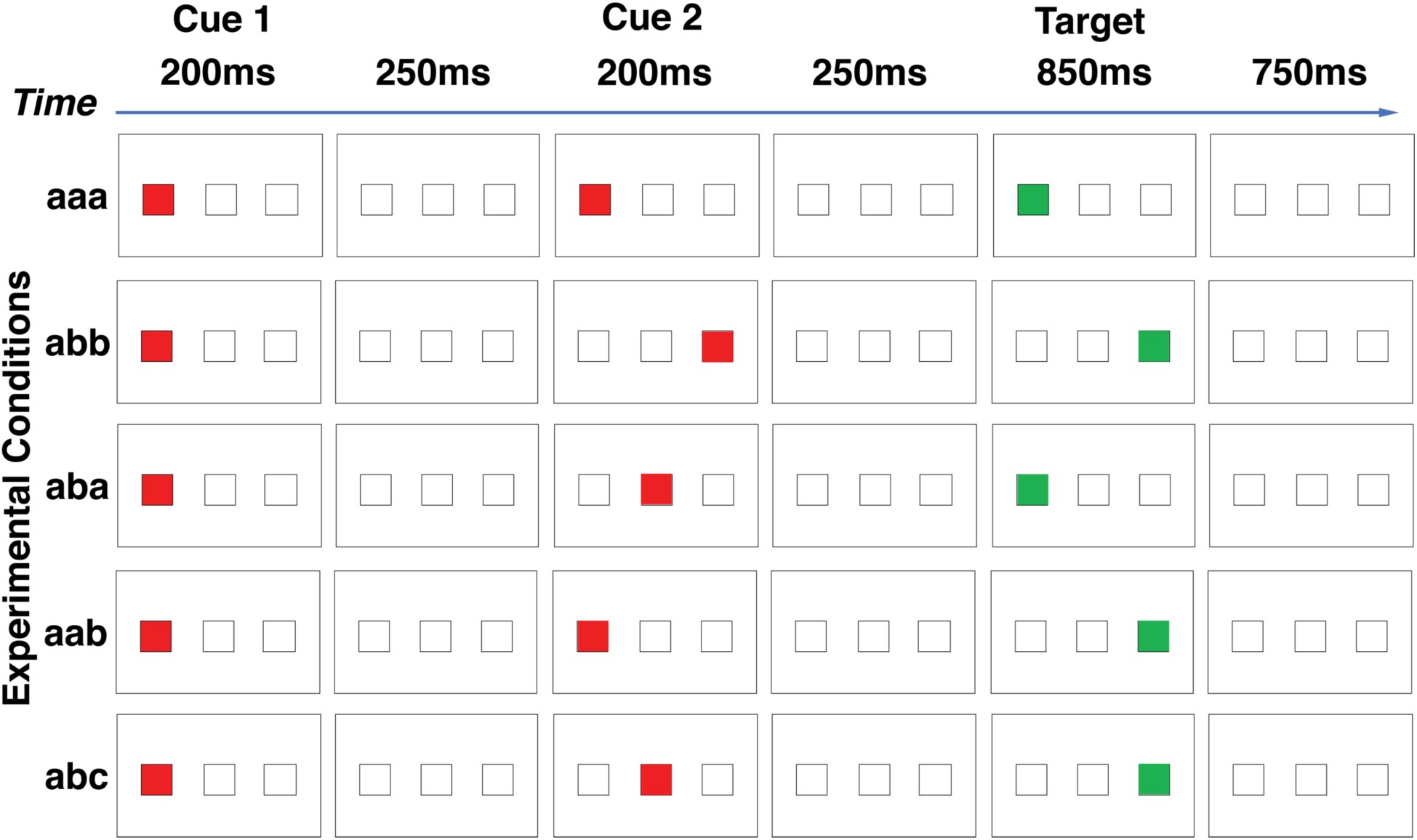
The double-cue spatial IOR experiment paradigm. Within each trial, there were three sequentially presented visual stimuli—two cues (solid red square) and one target (solid green square)—with a blank screen in between. The three stimuli were presented serially. The two cue stimuli could appear in any of the three locations (left, middle, right), whereas the target stimuli could only appear in one of the two locations (left or right, but not the middle). Subjects were instructed to respond to the target (solid green square) by pressing one of two buttons in the right hand to indicate whether the target was presented at the left or right location. The two cues were presented 200msec each, with a 250msec break in between. The second cue was followed by another 250msec break before the onset of the target, which was presented for 850msec. The next trial started 750msec after the offset of the target stimulus. There were five conditions based on the relationship of the locations in which the three stimuli were presented, *aaa, abb, aba, aab*, and *abc* (see main text). One example of each condition is shown here.

**Figure 2.**
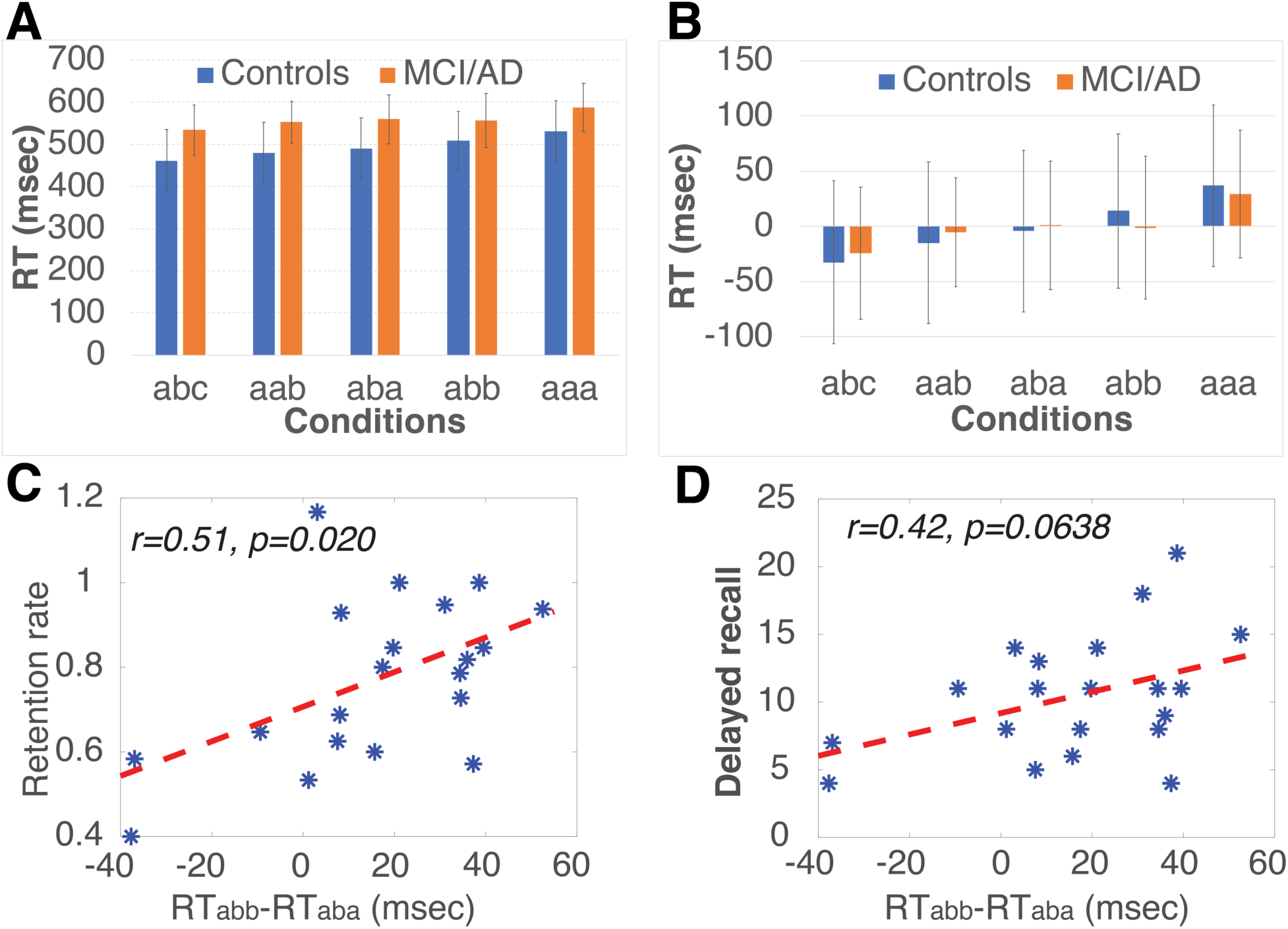
The double-cue spatial IOR experiment data. (A) Mean RT from the 16 MCI/AD patients and 41 age-matched controls. (B) The RT with mean RT removed from each subject for illustration purpose. (C, D) The correlation between spatial IOR (measured as the RT difference between two middle conditions, *abb* and *aba* (RT_abb_-RT_aba_)) and (D) the retention score (delayed recall divided by immediate recall) or (D) the delayed recall score in 20 high-risk controls (the subset of controls who have a family history of AD or at least one copy of APOE ε4 allele). Error bars represent standard deviation.

There were five conditions based on the locations in which the two cues and the target were sequentially presented (see Fig. 1A for an example of the five conditions).

***aaa***, in which the two cues and the target were presented at the same location;

***abb***, in which the second cue and the target were presented at the same location, and the first cue was presented at a different location;

***aba***, in which the first cue and the target were presented at the same location, and the second cue was presented at a different location;

***aab***, in which the two cues were presented at the same location, and the target was presented at a different location;

***abc***, in which the two cues and the target were presented at three different locations.

### Statistical Analysis

The group (controls vs MCI/AD) difference in demographics and neuropsychological tests were investigated using two-sample t-tests or Fisher’s Exact tests. The spatial IOR effects were investigated using mixed-design ANOVAs with one within-subject factor (experimental conditions, Fig. 1A), and one between-subject factor (controls vs MCI/AD). The F and p values of the coefficients in the mixed model ANOVA were adjusted using the Greenhouse-Geisser correction.

## RESULTS

The demographics, APOE ε4 status, and tests scores of the participants are shown in Table 1. A mean RT was determined for each condition and each participant. Mixed-design ANOVAs were carried out on these data.

A mixed-design ANOVA with a within-subject factor (Condition: *abc, aab, aba, abb, and aaa*) and a between-subject factor (Group: controls vs MCI/AD) revealed significant effects of Condition, F(4,216)=44.244, p<0.001, and Group, F(1,54)=10.295, p=0.002, as well as a trend for a Group x Condition interaction, F(4,216)=2.582, p=0.058 (Fig. 1B and 1C, Supplementary Table 1). To examine the spatial IOR impairment in MCI/AD patients further, we conducted two additional analyses.

First, a mixed-design ANOVA with the two “end” conditions (Condition: *abc* vs *aaa*) and a between-subject factor (Group: controls vs MCI/AD) revealed significant main effects of Condition, F(1,54)=99.827, p<0.001, and Group F(1,54)=10.148, p=0.002, but no significant interaction, F(1,54)=1.609, p=0.210.

Second, a mixed-design ANOVA on two “middle” conditions (Condition: *aba* vs *abb*) and the between-subject factor revealed significant main effects of Condition, F(1,54)=4.127, p=0.047, and Group, F(1,54)=8.328, p=0.006, as well as a significant Group x Condition interaction, F(1,54)=6.915, p=0.011. Taken together, the two additional analyses suggested that spatial IOR is impaired in individuals with MCI or mild AD, and the injury can be detected with the “more” sensitive comparison (i.e., conditions *aba* vs *abb*). The group difference between the two key conditions (*aba* and *abb*) was further confirmed by a permutation analysis (p_permutation_ = 0.00551), in which the group labels (controls vs MCI/AD) were randomly shuffled 100,000 times. In addition, the effect size of the group difference between the two key conditions (*aba* and *abb*) was 0.6325, suggesting a medium to large effect size.

Furthermore, we investigated whether the spatial IOR effects were sensitive to potential early AD pathological changes. In this analysis, we identified a subset of controls who had either a family history of AD or at least one APOE ε4 allele (“high-risk”, n=20). Then we investigated the relationship between the spatial IOR effect (measured as the difference between RT_*abb*_ and RT_*aba*_, Fig. 1B) and test scores in the “high-risk” group. Pearson’s correlation analyses revealed that the spatial IOR effect (RT_*abb*_ -RT_*aba*_) significantly correlated with retention rate (delayed-recall/immediate-recall, r=0.514, p=0.020, Fig. 1D), MMSE (r=0.520, p=0.019), LADL (r=0.676, p=0.001), and NPI (r=-0.480, p=0.032) scores, along with a trend with delayed recall (r=0.422, p=0.064, Fig. 1E).

## DISCUSSION

Previous studies using classic single cue-target paradigms report conflicting findings regarding spatial IOR impairment in individuals with MCI or AD. In the present study, we introduced a novel double cue paradigm capable of detecting the spatial IOR effect in a “finer” resolution (see the RT profiles of five conditions in Fig. 1B and 1C), observed spatial IOR impairment in individuals with MCI or mild AD. In addition, there was no difference in spatial IOR between individuals with MCI and individuals with mild AD, suggesting that spatial IOR impairment may occur at an early disease stage. This hypothesis was further supported by the data from the high-risk control subgroup, in which reduced spatial IOR effect correlated with reduced neuropsychological test scores, including delayed recall. The average age of those 20 high-risk controls was 66.0, suggesting that spatial IOR could be an early sign of underlying pathogenesis before the onset of cognitive impairment.

In AD, in addition to the medial temporal lobe (MTL), injury to the temporoparietal association cortex is frequently observed at early disease stages ^7,16^. For instance, amyloid plaques typically appear in the posterior association cortices prior to the MTL, and metabolic dysfunction is most frequently reported in the temporoparietal regions, which has a high accuracy in assisting AD diagnosis ^17^. Neural injury in the temporoparietal regions is also predictive of conversion from MCI to AD ^18^. Therefore, behavioral tests focused on temporoparietal and inferior parietal cortices may have the potential to assist MCI and early AD diagnosis, including the double cue spatial IOR task. Indeed, preliminary results using various machine learning algorithms suggest that integrating spatial IOR behavioral data with standard neuropsychological tests improves diagnostic accuracy ^19^. Furthermore, spatial IOR is robust and resistant to practice effect ^20,21^, thus making it an ideal tool in longitudinal studies or clinical trials.

There are some limitations of this study. The sample size limited our interpretation of the results, and large study could more thoroughly test those associations, especially the correlation between IOR effects and neurocognitive performance in “high-risk” controls. It would be interesting to investigate whether the difference between the conditions *aba* and *abb* is related to the difference in the IOR onset time ^22^. In addition, the neural mechanisms underlying the double cue spatial IOR impairment remain to be elucidated with multimodality neuroimaging studies, and longitudinal studies are required to examine whether the reduced spatial IOR effect in high-risk control individuals predicts progression to MCI or AD.

In conclusion, these findings support the notion that spatial IOR is impaired in individuals with MCI or mild AD, and the impairment is mild but detectable using the novel double cue paradigm. In addition, data from MCI/AD patients and high-risk controls suggest that spatial IOR impairment might occur at an early disease stage.

## Acknowledgements

We wish to thank all participants for their time and participation, Sikoya Ashburn for data collection, Amanda Dibattista for helping with APOE genotyping, Kyle Shattuck and Jessica Jacobs for their help with data analysis, the Memory Disorders Program at Georgetown University Medical Center, and the assistance for patient care from the Georgetown University Clinical Research Unit (GU-CRU), which has been funded in whole or in part with Federal funds (Grant # UL1TR000101 previously UL1RR031975) from the National Center for Advancing Translational Sciences (NCATS), National Institutes of Health (NIH), through the Clinical and Translational Science Awards Program (CTSA), a trademark of DHHS, part of the Roadmap Initiative, “Re-Engineering the Clinical Research Enterprise.”

The study was funded by the Alzheimer’s Drug Discovery Foundation (X.J.).

## Conflict of Interest

The authors declare no competing interests.

## Author contributions statements

XJ conceptualized the study. XJ and JHH designed the experiment. XJ and RST were involved in data collection. RST provided clinical diagnosis of MCI and AD patients. XJ analyzed the data. XJ wrote the manuscript. XJ, JHH, GWR, and RST revised the manuscript. All authors reviewed the final manuscript.

## Data Availability

The datasets are available upon reasonable request to the corresponding author.

## Figure Captions

**Supplementary Figure 1.**
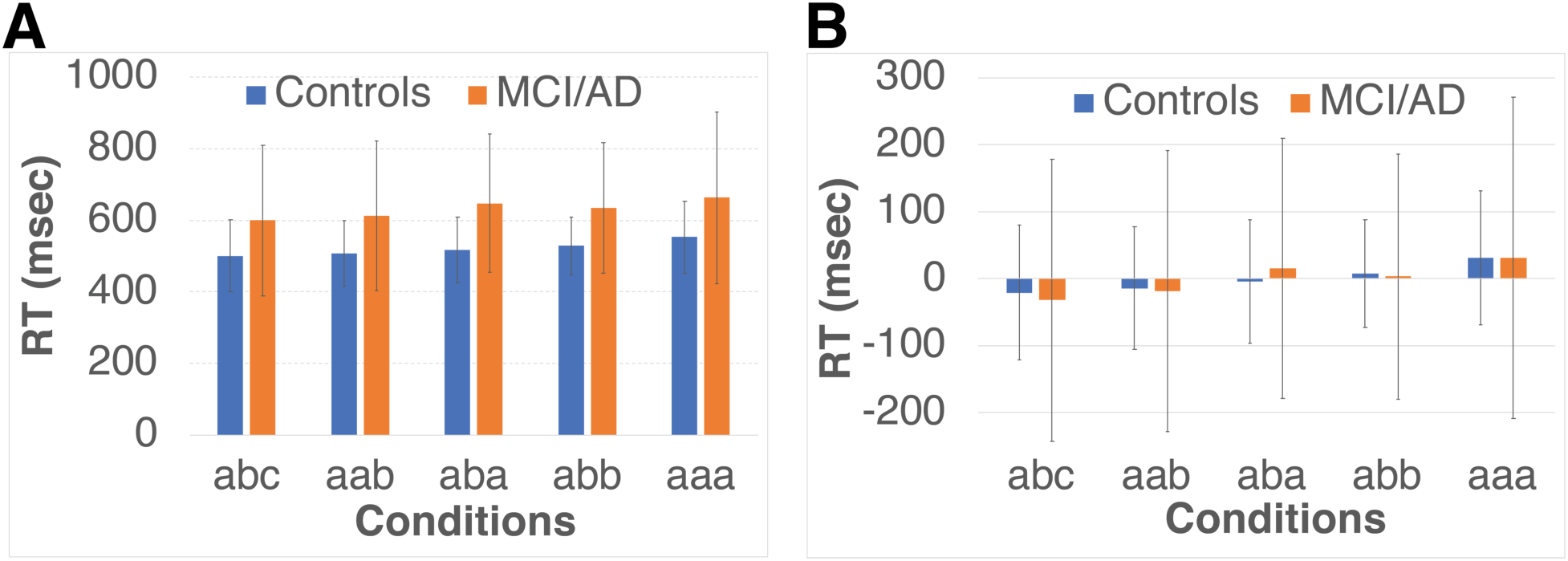
The double-cue spatial IOR data from the subjects who were excluded from the main analysis. Data from 9 AD, 4 MCI, and 9 control subjects were excluded from the data analysis in the main article due to younger than 58 (n=7), older than 80 (n=7), no high school diploma (n=2), with HIV-disease (n=1), or failed to perform the spatial IOR task (with an accuracy less than 75% due to failure to respond within the time window) (n=5). The data shown here include nine controls (age, 59.4±6.0 years old; 6 females) and 10 subjects with MCI or mild AD (age, 76.8±8.7 years old; 4 females; 3 MCI and 7 mild AD subjects). One MCI and two mild AD subjects did not have enough trials for each condition (with an overall accuracy less than 20%) to be included here. The data from these excluded subjects is in line with the data in the main analysis, suggesting a robust IOR deficit as measured by the difference between the conditions *aba* and *abb* (517ms and 529ms in the excluded controls, versus 647ms vs 635ms in the excluded MCI/mild AD subjects). Error bars represent standard deviation.

**Supplementary Figure 2.**
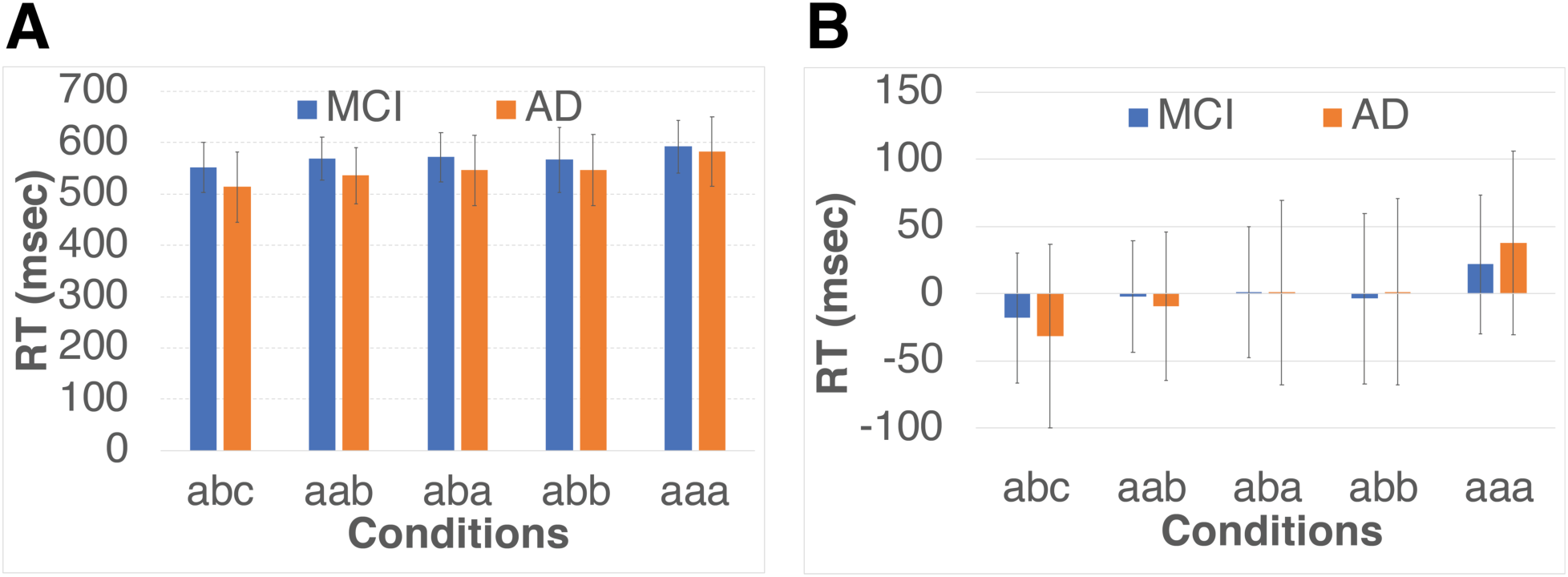
The double-cue spatial IOR data from the MCI (n=8; age, 71.3±3.1 years old; 2 females) and mild AD (n=7; age, 67.0±5.5 years old; 3 females) included in the main data analysis. A mixed-design ANOVA with a within-subject factor (Condition: *abc, aab, aba, abb, and aaa*) and a between-subject factor (Group: MCI vs AD) revealed significant effects of Condition, F(4,52)=10.301, p<0.001, but not of Group, F(1,13)=0.805, p=0.386, or the Group x Condition interaction, F(4,216)=0.855, p=0.476. Error bars represent standard deviation.

## Notes

### Competing Interest Statement

The authors have declared no competing interest.

## REFERENCES

1. Seidel Malkinson T, Bartolomeo P. Fronto-parietal organization for response times in inhibition of return: The FORTIOR model. Cortex J Devoted Study Nerv Syst Behav. 2018;102:176–192. doi:10.1016/j.cortex.2017.11.005

2. Posner MI, Cohen, Y. Components of visual orienting. In: Attention and Performance X. edited by Bouma, H.; and Bouwhuis, D.G. Hillsdale (NJ): Lawrence Erlbaum; 1984:531–556.

3. Spence C, Lloyd D, McGlone F, Nicholls ME, Driver J. Inhibition of return is supramodal: a demonstration between all possible pairings of vision, touch, and audition. Exp Brain Res. 2000;134(1):42–48. doi:10.1007/s002210000442

4. Briand KA, Larrison AL, Sereno AB. Inhibition of return in manual and saccadic response systems. Percept Psychophys. 2000;62(8):1512–1524. doi:10.3758/bf03212152

5. MacInnes WJ. Multiple Diffusion Models to Compare Saccadic and Manual Responses for Inhibition of Return. Neural Comput. 2017;29(3):804–824. doi:10.1162/NECO_a_00904

6. Hartley AA, Kieley JM. Adult age differences in the inhibition of return of visual attention. Psychol Aging. 1995;10(4):670–683.

7. Besson FL, La Joie R, Doeuvre L, et al. Cognitive and Brain Profiles Associated with Current Neuroimaging Biomarkers of Preclinical Alzheimer’s Disease. J Neurosci Off J Soc Neurosci. 2015;35(29):10402–10411. doi:10.1523/JNEUROSCI.0150-15.2015

8. Amieva H, Phillips LH, Della Sala S, Henry JD. Inhibitory functioning in Alzheimer’s disease. Brain J Neurol. 2004;127(Pt 5):949–964. doi:10.1093/brain/awh045

9. Faust ME, Balota DA. Inhibition of return and visuospatial attention in healthy older adults and individuals with dementia of the Alzheimer type. Neuropsychology. 1997;11(1):13–29.

10. Danckert J, Maruff P, Crowe S, Currie J. Inhibitory processes in covert orienting in patients with Alzheimer’s disease. Neuropsychology. 1998;12(2):225–241.

11. Tales A, Snowden RJ, Phillips M, et al. Exogenous phasic alerting and spatial orienting in mild cognitive impairment compared to healthy ageing: Study outcome is related to target response. Cortex J Devoted Study Nerv Syst Behav. 2011;47(2):180–190. doi:10.1016/j.cortex.2009.09.007

12. Tales A, Snowden RJ, Haworth J, Wilcock G. Abnormal spatial and non-spatial cueing effects in mild cognitive impairment and Alzheimer’s disease. Neurocase. 2005;11(1):85–92. doi:10.1080/13554790490896983

13. Bayer A, Phillips M, Porter G, Leonards U, Bompas A, Tales A. Abnormal inhibition of return in mild cognitive impairment: is it specific to the presence of prodromal dementia? J Alzheimers Dis JAD. 2014;40(1):177–189. doi:10.3233/JAD-131934

14. Dukewich KR, Boehnke SE. Cue repetition increases inhibition of return. Neurosci Lett. 2008;448(3):231–235. doi:10.1016/j.neulet.2008.10.063

15. Howard JH, Howard DV, Dennis NA, Kelly AJ. Implicit learning of predictive relationships in three-element visual sequences by young and old adults. J Exp Psychol Learn Mem Cogn. 2008;34(5):1139–1157. doi:10.1037/a0012797

16. Chiotis K, Saint-Aubert L, Rodriguez-Vieitez E, et al. Longitudinal changes of tau PET imaging in relation to hypometabolism in prodromal and Alzheimer’s disease dementia. Mol Psychiatry. 2018;23(7):1666–1673. doi:10.1038/mp.2017.108

17. Sperling RA, Aisen PS, Beckett LA, et al. Toward defining the preclinical stages of Alzheimer’s disease: recommendations from the National Institute on Aging-Alzheimer’s Association workgroups on diagnostic guidelines for Alzheimer’s disease. Alzheimers Dement J Alzheimers Assoc. 2011;7(3):280–292. doi:10.1016/j.jalz.2011.03.003

18. Pagani M, Nobili F, Morbelli S, et al. Early identification of MCI converting to AD: a FDG PET study. Eur J Nucl Med Mol Imaging. 2017;44(12):2042–2052. doi:10.1007/s00259-017-3761-x

19. Almubark I, Chang L-C, Nguyen T, Turner RS, Jiang X. Early Detection of Alzheimer’s Disease Using Patient Neuropsychological and Cognitive Data and Machine Learning Techniques. In: The IEEE BigData 2019 Conference.; 2019:Poster #276. http://bigdataieee.org/BigData2019/files/BigData2019PosterList.pdf.

20. Bao Y, Sander T, Trahms L, Pöppel E, Lei Q, Zhou B. The eccentricity effect of inhibition of return is resistant to practice. Neurosci Lett. 2011;500(1):47–51. doi:10.1016/j.neulet.2011.06.003

21. Pratt J, McAuliffe J. Examining the effect of practice on inhibition of return in static displays. Percept Psychophys. 1999;61(4):756–765. doi:10.3758/bf03205543

22. Li T, Wang L, Huang W, et al. Onset Time of Inhibition of Return Is a Promising Index for Assessing Cognitive Functions in Older Adults. J Gerontol B Psychol Sci Soc Sci. 2020;75(4):753–761. doi:10.1093/geronb/gby070

